# Targeting F-actin stress fibers to suppress the dedifferentiated phenotype in chondrocytes

**DOI:** 10.1101/2023.12.08.570865

**Authors:** Mandy M. Schofield, Alissa Rzepski, Jonah Hammerstedt, Sohan Shah, Chloe Mirack, Justin Parreno

## Abstract

Actin is a central mediator of the chondrocyte phenotype. Monolayer expansion of articular chondrocytes on tissue culture polystyrene, for cell-based repair therapies, leads to chondrocyte dedifferentiation. During dedifferentiation, chondrocytes spread and filamentous (F-)actin reorganizes from a cortical to a stress fiber arrangement causing a reduction in cartilage matrix expression and an increase in fibroblastic matrix and contractile molecule expression. While the downstream mechanisms regulating chondrocyte molecular expression by alterations in F-actin organization have become elucidated, the critical upstream regulators of F-actin networks in chondrocytes are not completely known. Tropomyosin (TPM) and the RhoGTPases are known regulators of F-actin networks. The purpose of this study is to elucidate the regulation of passaged chondrocyte F-actin stress fiber networks and cell phenotype by the specific TPM, TPM3.1, and the RhoGTPase, CDC42. Our results demonstrated that TPM3.1 associates with cortical F-actin and stress fiber F-actin in primary and passaged chondrocytes, respectively. In passaged cells, we found that TPM3.1 inhibition causes F-actin reorganization from stress fibers back to cortical F-actin and also causes an increase in G/F-actin. CDC42 inhibition also causes formation of cortical F-actin. However, CDC42 inhibition, but not TPM3.1 inhibition, leads to the re-association of TPM3.1 with cortical F-actin. Both TPM3.1 and CDC42 inhibition reduces nuclear localization of myocardin related transcription factor, which is known to suppress dedifferentiated molecule expression. We confirmed that TPM3.1 or CDC42 inhibition partially redifferentiates passaged cells by reducing fibroblast matrix and contractile expression, and increasing chondrogenic SOX9 expression. A further understanding on the regulation of F-actin in passaged cells may lead into new insights to stimulate cartilage matrix expression in cells for regenerative therapies.

## Introduction

Chondrocyte dedifferentiation is the process by which cartilage cells undergo phenotypic alterations exemplified by a decrease in expression for cartilage matrix molecules and an increase in fibroblastic molecules (Abbott and Holtzer, 1968; Benya et al., 1978; Charlier et al., 2019; Gay et al., 1976; Mallein-Gerin et al., 1991; Parreno et al., 2017a). While chondrocyte dedifferentiation is thought to occur *in vivo* during disease progression in Osteoarthritis, dedifferentiation has been well characterized during monolayer expansion of chondrocytes for cell-based therapies (i.e. autologous chondrocyte implantation) and cartilage bioengineering (Bianchi et al., 2019; Kwon et al., 2019; Makris et al., 2015; Malicev et al., 2009; Parreno et al., 2018; Taylor et al., 2015). As compared to freshly isolated primary (P0) chondrocytes, chondrocytes that have been expanded in monolayer culture (passaged cells) are larger and more elongated. Rather than expressing cartilage matrix molecules, such as collagen type II (COL2) and aggrecan (ACAN), passaged cells produce fibroblast matrix molecules, such as Type I Collagen (COL1) and Tenascin C (TNC) (Benya et al., 1978; Holtzer et al., 1960; Mallein-Gerin et al., 1991; Parreno et al., 2017a; Parreno et al., 2014). Additionally, passaged cells elevate expression for contractile molecules α-smooth muscle actin (αSMA) and transgelin (TAGLN) (Kinner and Spector, 2001; Parreno et al., 2017b). Dedifferentiation is problematic for cell-based therapies as the tissue formed by passaged cells is fibrocartilage, which is biomechanically inferior to articular cartilage (Kinner and Spector, 2001; Mesallati et al., 2014; Murphy et al., 2015; Roberts et al., 2009). The increase in cellular contractility can also lead to matrix contraction and failure of repair tissue to integrate with native tissue (Lee et al., 2003; Parreno et al., 2017b). Therefore, understanding the regulatory mechanisms involved in dedifferentiation may lead to new targets to improve cartilage repair therapies.

The actin cytoskeleton is a potent regulator of the dedifferentiated phenotype (Benya et al., 1988; Mallein-Gerin et al., 1991; Parreno et al., 2017a). In primary chondrocytes, filamentous (F-)actin is cortically arranged. However, monolayer expansion results in the reorganization of cortical F-actin into stress fibers (Parreno et al., 2014). In addition, the formation of F-actin stress fibers coincides with a change in actin polymerization status. As compared to primary chondrocytes which have a high proportion of globular (G-) to F-actin (G/F-actin), passaged chondrocytes have reduced proportion of G/F-actin (Parreno et al., 2017a; Parreno et al., 2014; Parreno et al., 2017b). Preventing cell spreading by placing cells on low/non-adherent substrates or treating with compounds that prevent polymerization, such as latrunculins or cytochalasins, eliminate actin stress fibers, increase G/F-actin, leading to increased chondrogenic molecules SRY-Box Transcription Factor 9 (SOX9) and ACAN expression, as well as decreasing fibroblastic and contractile markers (Parreno et al., 2014; Parreno et al., 2017b).

Actin polymerization directly regulates gene expression via a G-actin binding transcription factor, myocardin-related transcription factor-a (MRTF) (Asparuhova et al., 2011; Luchsinger et al., 2011). The binding of MRTF to G-actin sequesters MRTF to the cytoplasm of cells (Kuwahara et al., 2005). When G-actin polymerizes into F-actin, MRTF is liberated from G-actin and translocates into the nucleus of cells (Kuwahara et al., 2005). In the nucleus, MRTF enhances the expression of fibroblast matrix and contractile genes through its interaction with serum response factor (Esnault et al., 2014; Parmacek, 2007). In passaged chondrocytes, where G/F-actin is relatively low as compared to primary chondrocytes, MRTF exists predominantly in the nucleus of cells. MRTF inhibition or knockdown reduces fibroblast matrix (*COL1* and *TNC*) as well as contractile molecules α*SMA* and *TAGLN* (Parreno et al., 2014; Parreno et al., 2017b).

Targeting the actin cytoskeleton in passaged chondrocytes is a means to repress dedifferentiation. While preventing actin polymerization by exposure of cells to latrunculin or cytochalasin may lead to favorable effects (Delve et al., 2018; Parreno et al., 2014; Parreno et al., 2017b), such compounds do not target specific F-actin networks. These drugs have been shown to prevent the formation of cortical F-actin as well as can lead to cell death (Park et al., 2008; Parreno et al., 2017a; Parreno et al., 2014; Parreno et al., 2017b). Thus, approaches to promote passaged chondrocytes redifferentiation by specifically targeting F-actin stress fibers while allowing for cortical actin organization are required. In this study, we examine the potential to regulate F-actin organization and cellular phenotype in passaged chondrocytes by targeting two known regulators of actin, the Tropomyosins (TPMs) and RhoGTPases.

TPMs are master regulators of F-actin organization (Gunning et al., 2015). They are long, rod-like proteins that bind and stabilize F-actin. TPMs also determine F-actin network reorganization by facilitating the interaction of F-actin with other actin binding proteins, including actin severing, capping, contractile, cross- linking, and depolymerizing proteins (Bryce et al., 2003; Gunning et al., 2015; Schevzov et al., 2011). TPMs bind along F-actin and can either interfere or promote the binding of specific actin binding proteins with F-actin (Maekawa and Toriyama, 1994). The expression of specific TPM isoforms by a cell type provides combinatorial diversity to define F-actin organization. In particular, the TPM isoform, TPM3.1, has been shown to associate with and stabilize F-actin in stress fibers in lens epithelial cells (Parreno et al., 2020), tenocytes (Inguito et al., 2022) and U2OS cells (Gateva et al., 2017). However, it remains to be determined if TPM3.1 stabilizes F-actin stress fibers in passaged chondrocytes. If so, targeting TPM3.1 could provide a way to repress F-actin stress fibers and suppress dedifferentiation for cell-based therapies or bioengineering purposes.

The RhoGTPases are a family of molecular switches that regulate the F-actin cytoskeleton and cellular shape (Ridley and Hall, 1992). The three major Rho-GTPases are: (1) Ras-related guanosine triphosphate (GTP)-binding proteins Ras homolog gene family, member A (RHOA); (2) Ras-related C3 botulinum toxin substrate 1 (RAC); and, (3) cell division control protein 42 (CDC42). RHO, RAC, and CDC42 are inactive when bound to guanosine diphosphate (GDP). Guanine exchange factor (GEF) promotes the dissociation of GDP from GTPase allowing GTP to bind and interact with effector, actin binding proteins (Kohno et al., 1996; Kurisu and Takenawa, 2009). All three Rho-GTPases have been shown to regulate stress fiber formation in cells (Lauer et al., 2021). In chondrocytes, monolayer expansion increases the expression of RhoGTPases RHOA and RAC1, and CDC42 (Shin et al., 2016). While these findings would suggest that RhoGTPase inhibition may be a means to suppress the dedifferentiated phenotype, RhoGTPase inhibition studies have led to conflicting results. Inhibition of RhoA signaling by targeting downstream effector, Rho-associated kinase (ROCK), inhibitor suppress the chondrocyte dedifferentiation in several studies (Matsumoto et al., 2012; Tew and Hardingham, 2006; Woods et al., 2005), but another study has shown little to no effect by inhibition of ROCK or RAC1 (Shin et al., 2016). With regard to CDC42, it has been shown that expression of dominant negative CDC42 decreases F-actin stress fibers in chondrocytes leading to an increase in chondrogenic expression (Fortier et al., 2004). However, it has also been demonstrated that treatment of primary chondrocytes with inflammatory cytokines (IL-1, IL-6, IL-8) unexpectedly decreases active GTP-CDC42 but still promotes F-actin stress fiber formation (Novakofski et al., 2012). Therefore, the role that CDC42 plays in regulating stress fiber formation in monolayer expanded chondrocytes is unclear. Further investigation on the regulation of the differentiated phenotype by RhoGTPases is required.

Here, we investigated the regulation of cell shape, F-actin organization and downstream actin based signaling by TPM3.1 or the RhoGTPases. We test the hypothesis that by promoting cell rounding and reducing F-actin, through targeting TPM3.1 or specific RhoGTPases, will repress MRTF localization and the expression of dedifferentiated molecules.

## Methods

### Cell Isolation and Culture

Bovine cells were isolated and cultured as previously described (Parreno et al., 2014), with slight modifications. Briefly, cartilage was harvested from bovine metacarpophalangeal joints that were obtained from a local butcher within 24 hours of euthanasia. Cartilage pieces were placed in 0.5% proteinase (Sigma-Aldrich; Burlington, MA) at 37°C. After 1 hour, cartilage was washed three times in phosphate buffered saline (PBS; potassium phosphate, sodium chloride, sodium phosphate dibasic; GenClone; Genesee Scientific, San Diego, CA, USA) and then placed in 0.1% collagenase (Roche; Burlington, MA). After an overnight digestion at 37°C, cell suspensions were passed through a 100micron cell strainer (GenClone). Cells were then pelleted by centrifugation at 600g for 8 minutes, resuspended in expansion media consisting of Ham’s F-12 (Corning; Manassas, VA) supplemented with 10% Fetal Bovine Serum (FBS) (GenClone) and 1% antibiotic-antimycotic (MilliporeSigma). Cells were seeded on plastic tissue culture flasks at a concentration of 2.0x10^3^ cells/cm^2^. Culture media was replenished every 2-3 days. Once cells reached 70-80% confluency, they were detached from culture flasks using 0.25% Trypsin (GenClone). At this point they were regarded as passage 1 (P1) cells. P1 cells were pelleted and then re-seeded at 2.0 x 10^3^ cells/cm^2^. Cells remained on culture vessels until they reached 70-80% confluency at which point they were detached from culture vessels and deemed as passage 2 (P2) cells.

Human cells were cultured as previously described (Parreno et al., 2017b) with slight modifications. Human P0 cells were obtained from StemBioSys (San Antonio, TX). Human P0 cells were seeded on plastic tissue culture flasks at a concentration of 2000 cells/cm^2^. Culture media consisted of high glucose Dulbecco’s Modified Eagle’s Medium (DMEM) (GenClone, Genesee Scientific, San Diego, CA, USA) supplemented with 20% Fetal Bovine Serum (FBS) (GenClone, Genesee Scientific; San Diego, CA, USA) and 1% antibiotic-antimycotic (MilliporeSigma) and was replenished every 2-3 days. Once cells reached 70-80% confluency, they were detached and passaged to P2 in the same manner as bovine cells.

### Pharmacological Inhibition

For TPM3.1 inhibition, P2 cells were seeded for experiments at a concentration of 2.6x10^4^ cells/cm^2^ in expansion media. After 24 hours, cells were exposed to TPM3.1 inhibitors TR100 (MilliporeSigma) or ATM3507 (MedChemExpress; Monmouth Junction, NJ). Preliminary experiments indicated that exposure of cells to 8μM TR100 and ATM3507 began to lead to cell death. Therefore, for our studies, we used concentrations below 8μM. For TR100, the effective TR100 concentrations of 4μM and 6μM for bovine and human cells, respectively, were used. For ATM3507, we used the effective concentration of 6μM for both bovine and human cells. DMSO (ThermoFisher Scientific; Waltham, MA) was used as a vehicle control.

To inhibit RhoGTPases, P2 cells were seeded for experiments at a concentration of 2.6x10^4^ cells/cm^2^ in serum starved (0.5%FBS) Ham’s F12 with effective concentrations of RhoGTPase inhibitors: RAC inhibitor NSC23766 (50 μM), ROCK inhibitor Y27632 (10 μM) (Tocris; Minneapolis, MN, USA), CDC42 inhibitor ML141 (10 μM) (MilliporeSigma). DMSO was used as a vehicle for all the inhibitors used in this study.

### Light Microscopy and cell area and circularity quantification

Following 24 hours of inhibitor treatment, cells were imaged on a Axiovert 25 inverted phase-contrast microscope (Zeiss; Jena, Germany) using an attached camera (Swiftcam Technologies; Hong Kong, Japan). The boundaries of individual cells were manually traced on images using FIJI software. Cell area and circularity were then calculated. Circularity was defined as C = 4π (A/P^2^) where, P is the perimeter, and A is the cell area. A circularity value of 0 indicates an elongated ellipse, whereas a value of 1 indicates a perfect circle.

Live cells were additionally imaged using CytoSMART Lux2 live cell analysis platform (CytoSMART; Eindhoven, The Netherlands) for 24 hours with images being captured every 15 minutes. Cell tracings were performed at 0, 12, and 24 hours and morphology was analyzed.

### RNA Extraction, Reverse Transcription, and Semi-Quantitative / Relative Polymerase Chain Reaction

RNA was isolated from cells using TRIzol (Sigma-Aldrich) as previously described (Inguito et al., 2022). Phase separation was performed using chloroform, followed by precipitation in isopropanol. RNA precipitates were washed in ethanol, briefly air-dried and then resuspended in molecular grade water (IBI Scientific; Dubuque, IA). RNA was reverse transcribed to cDNA with the UltraScript 2.0 cDNA Synthesis Kit (PCR Biosystems; Wayne, PA).

Semi-quantitative gel electrophoresis PCR was performed on equal concentrations of cDNA using 2x PCRBIO HS Taq Mix Red (PCR Biosystems; London, UK) per manufacturer’s instructions on a Sapphire Biomolecular Imager (Azure; Houston, TX, USA). cDNA samples plus 2x PCRBIO HS Taq Mix Red (PCR Biosystems) underwent 40 PCR cycles in a Thermocycler (Hercules, CA) before undergoing gel electrophoresis. To perform Sanger sequencing of TPM3.1, gel electrophoresis was performed on PCR products. Using a UVP UV Transilluminator (Upland, CA), bands corresponding to the product size of TPM3.1 in the gel were cut out using a sharp blade and sent for Sanger Sequencing (Azenta Life Sciences; Genewiz, South Plainfield, NJ).

Relative real-time RT-PCR was performed using equal concentrations of cDNA, with qPCRBio Sygreen Blue Mix (PCR Biosystems), as per manufacturer’s directions on a Cielo 3 real-time PCR machine (Azure; Houston, TX, USA). Primers used are listed in Table 1. mRNA levels were calculated using a ΔΔCT method (Schmittgen and Livak, 2008) with normalization to *18S*.

**Table 1.**
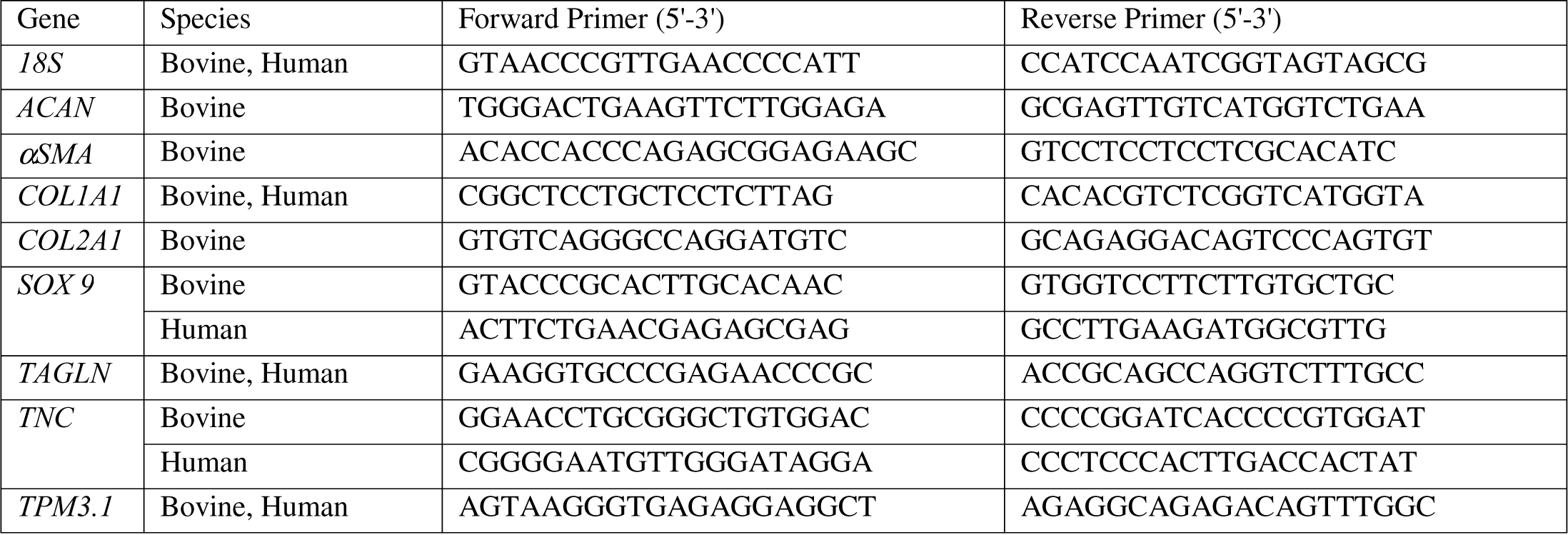
PCR primers used in Study.

### WES Capillary Electrophoresis

Protein levels in cells were analyzed as previously described (Parreno et al., 2022). Briefly, cells were seeded, and removed from culture vessels via cell scraping and centrifuged at 800g for 8 minutes. Protein was extracted by resuspending cell pellets in Radioimmunoprecipitation (RIPA) lysis buffer Millipore Sigma), sonicated (Model Q55, Qsonica; Newton, CT) and quantified using a bicinchoninic acid (BCA) protein assay (Prometheus Protein Biology Products; Genesee Scientific, San Diego, CA, USA). Protein levels were quantified using WES capillary electrophoresis (Protein Simple, San Jose, CA, USA) according to manufacturer’s instructions. Protein lysates (0.5 μg/mL) was loaded per well of either a 12-230kDa or 66- 440kDa separation module plate with antibodies listed in Table 2.

**Table 2.** Primary Antibodies used in Study.

| Target | Application | Species Isotype | Dilution | Company; Cat # |
| --- | --- | --- | --- | --- |
| Pan Actin | WES Capillary Electrophoresis | Rabbit polyclonal IgG | 1:100 | Cell Signaling; 4968 |
| αSMA | WES Capillary Electrophoresis | Mouse polyclonal IgG | 1:50 | Abcam; ab7817 |
| COL1α1 | WES Capillary Electrophoresis | Rabbit polyclonal IgG | 1:200 | Novus; NBP1-30054 |
| MRTF | Immunocytochemistry | Rabbit monoclonal IgG | 1:200 | Cell Signaling; 77098 |
| SOX9 | WES Capillary Electrophoresis | Rabbit polyclonal IgG | 1:10 | Novus; NBP2-24659 |
| TAGLN | WES Capillary Electrophoresis | Rabbit polyclonal IgG | 1:100 | Abcam; ab14106 |
| TPM3.1 | Immunocytochemistry | Mouse monoclonal IgG | 1:200 | Millipore Sigma; 2G10.2 |

### Triton Solubility Assay

Actin polymerization status was determined via Triton solubility assay using a previously published protocol with slight modifications (Parreno et al., 2017a; Parreno et al., 2014). To isolate soluble portions (containing predominantly G-actin), cells were placed in 0.1 %Triton in cytoskeletal buffer (100 mM NaCl, 3 mM MgCl2, 300 mM Sucrose, 1 mM EGTA, 10 mM PIPES). Solution was removed from dishes and then placed in microcentrifuge tubes. RIPA (10x) was added to the Triton soluble portions and placed on ice. To isolate the remaining cytoskeletal Triton insoluble portions (containing predominantly F-actin), the cytoskeletal fraction of cells that remained on culture dishes were placed in 0.1%Triton in cytoskeletal buffer containing 1X RIPA. The cell contents were scraped from dishes using cell scrapers. Samples were incubated on ice for at least 30 minutes and then sonicated. Equal volumes of Triton soluble and insoluble portions were prepared for WES capillary electrophoresis and separated on a 12-230 kDa separation module (Protein Simple, San Jose; CA, USA) using a Pan-actin antibody (Table 2).

### Confocal Microscopy and Quantification

Cells seeded on glass dishes were fixed using 4% paraformaldehyde (PFA) in PBS. Cells were permeabilized using permeabilization/blocking buffer consisting of 3% Goat serum, 3% BSA, and 0.3% Triton with primary antibodies overnight at 4°C. After three 5-minute washes, secondary antibody solution which contained AlexaFluor488 (1:200, Thermo Fisher Scientific), Rhodamine phalloidin (1:50; Biotium; Fremont, CA), and Hoechst 33342 (1:500; Biotium) to visualize specific proteins (i.e. TPM3.1 or MRTF), F-actin, and nuclei respectively were applied for 1 hour at room temperature. Stained cells were then mounted using ProLong gold anti-fade reagent (Thermo Fisher Scientific) and imaged on a LSM880 confocal microscope. Confocal images were processed using ZEN Blue (Zeiss).

### Image Analysis and Quantification

Ratiometric nuclear to cytoplasmic ratio analysis (nuclear:cytoplasmic) of MRTF fluorescence was performed using FIJI. MRTF fluorescence in the nucleus and cytoplasm was measured at the mid-region of the cells, where nuclear staining for Hoescht was brightest. Rhodamine phalloidin staining at the cell borders was used as an indicator to trace the cellular boundaries. The raw integrated densities of MRTF signal and area were obtained for the individual cell and the nucleus and used to calculate nuclear:cytoplasmic signal.

### Statistical Analysis

Experiments were conducted on at least 3 separate occasions, on samples from separate individuals or animals. Statistical analysis was performed using Graphpad Prism 9 (San Diego, CA, USA). Pooled data points from experiments were evaluated for outliers using the ROUT method with a maximum desired false discovery rate (Q) set at 1% (Motulsky and Brown, 2006). Differences between two groups of data were analyzed using unpaired t-tests. Differences between three or more groups were identified using an analysis of variance (ANOVA) followed by Dunnett post hoc test.

## RESULTS

*TPM3.1 is expressed and associates with F-actin in chondrocytes*

To determine if chondrocytes express *TPM*3.1, we performed reverse transcriptase-PCR (RT-PCR) for full-length *TPM3*.1 on bovine and human P2 cell cDNA (Figure S1). Gel electrophoresis shows bands at the expected sizes of 970bp for bovine and human *TPM3*.1, respectively. Sanger sequencing of isolated DNA from bands confirmed PCR products to be *TPM3.1* cDNA.

We next sought to determine protein localization of TPM3.1 in primary and passaged cells. Cells were immunostained for TPM3.1 using the 2G10.2 antibody which recognizes the 9D exon of TPM3.1. F-actin is cortically arranged in primary bovine chondrocytes (Figure 1A, B). Immunostaining for TPM3.1 reveals that TPM3.1 is also cortically arranged and appears to associate with cortical F-actin (Figure 1A,-C). Line scan analysis of F-actin and TPM3.1 fluorescent intensity supports this association as there is strong overlap between F-actin and TPM3.1 fluorescent peaks (Figure 1D). High magnification Airyscan imaging demonstrates TPM3.1 is associated with F-actin in microdomains (Figure 1C). In both passaged bovine and human cells, F-actin is organized into stress fibers (Figure 1A, B). TPM3.1 associates with F-actin stress fibers in both bovine and human passaged cells which is supported by line scan analysis of TPM3.1 and F-actin fluorescent intensity (Figure 1D). Similar to primary bovine chondrocytes, TPM3.1 organizes into microdomains along F-actin (Figure 1C).

**Figure 1.**
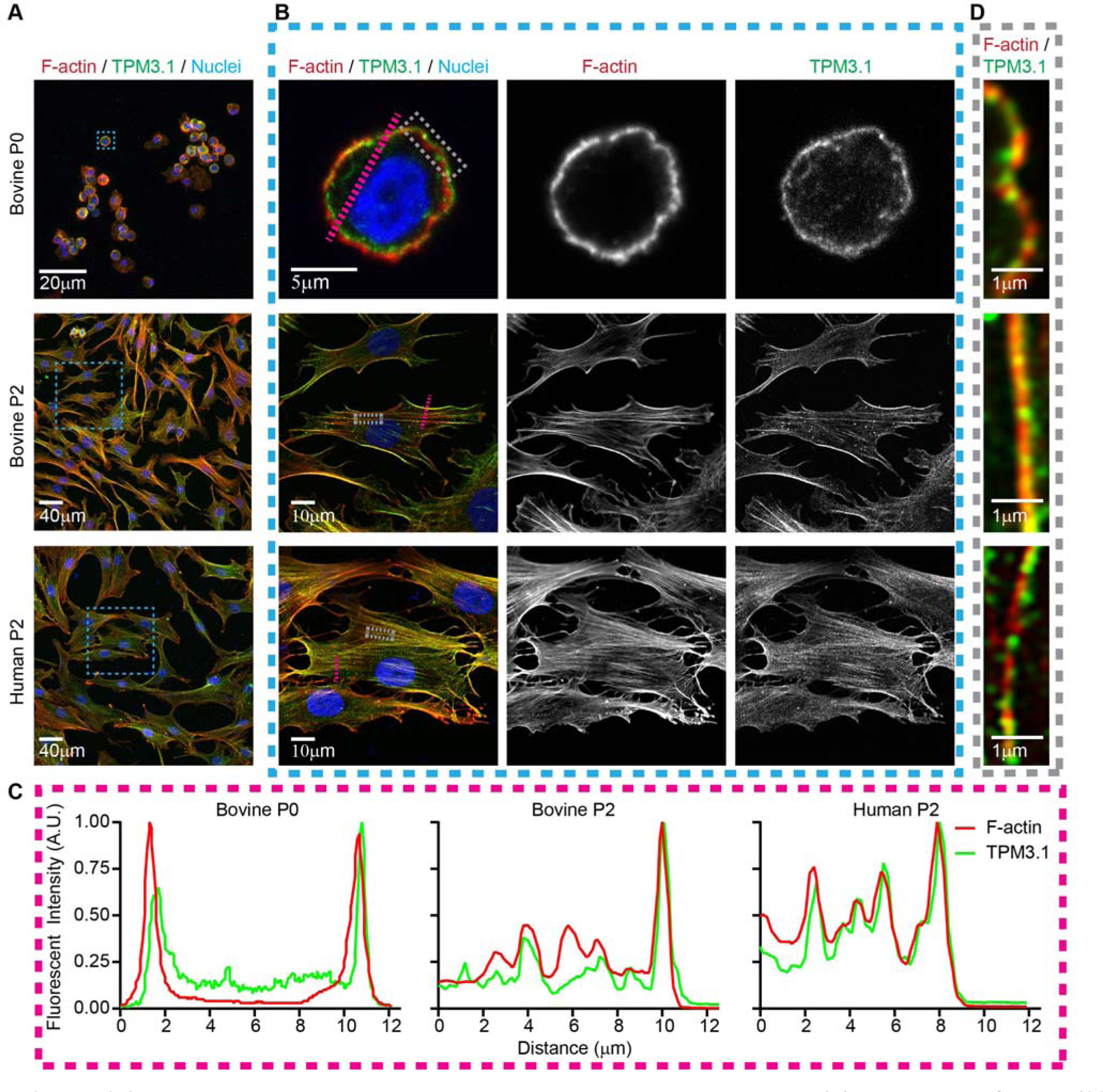
Tpm3.1 associates with F-actin in bovine and human chondrocytes. (A) Lower magnification (20x objective) maximum intensity projection of cells. (B) Higher magnification (63x objective) single optical section images of cells (blue dashed boxes) in panel A showing association of Tpm3.1 with cortical (P0 chondrocytes) and stress fiber F-actin (P2 chondrocytes). (C) Zoomed in Airyscan high magnification (63x objective) image of regions (grey dashed boxes) in panel B shows that Tpm3.1 is organized in microdomains along F-actin. Cells stained for F-actin (Phalloidin; Red), Tpm3.1 (Antibody 2G10.2; Green), and Nuclei (Hoechst; Blue). (D) Line scan analysis of lines (pink dashed) in panel B, demonstrating colocalization of Tpm3.1 with F-actin.

*TPM3.1 inhibition alters cell morphology and F-actin organization in passaged cells*

To evaluate the role of TPM3.1 in regulating passaged cells dedifferentiated phenotype, we treated bovine passaged chondrocytes with known TPM3.1 inhibitors, TR100 or ATM3507. These inhibitors are structurally diverse incorporate into the head-to-tail overlap of TPM3.1. When present during co-polymerization of TPM3.1 with actin, TR100 prevents TPM3.1 dependent F-actin polymerization. Extended exposure of the drug can lead to the breakdown of TPM3.1 bound actin filaments (Stehn et al., 2013).

We determined that TPM3.1 inhibition, by exposure of P2 cells to either TR100 and ATM3507, decreases cell area in bovine and human passaged cells (Fig. 2A, B). Additionally, TPM3.1 inhibition increases average cell circularity. Despite the increase in average cell circularity by TPM3.1 inhibition, there is a subpopulation of cells that remain elongated, in both passaged bovine and human chondrocytes as indicated by proportion of cells with low circularity values (Figure 2B).

**Figure 2.**
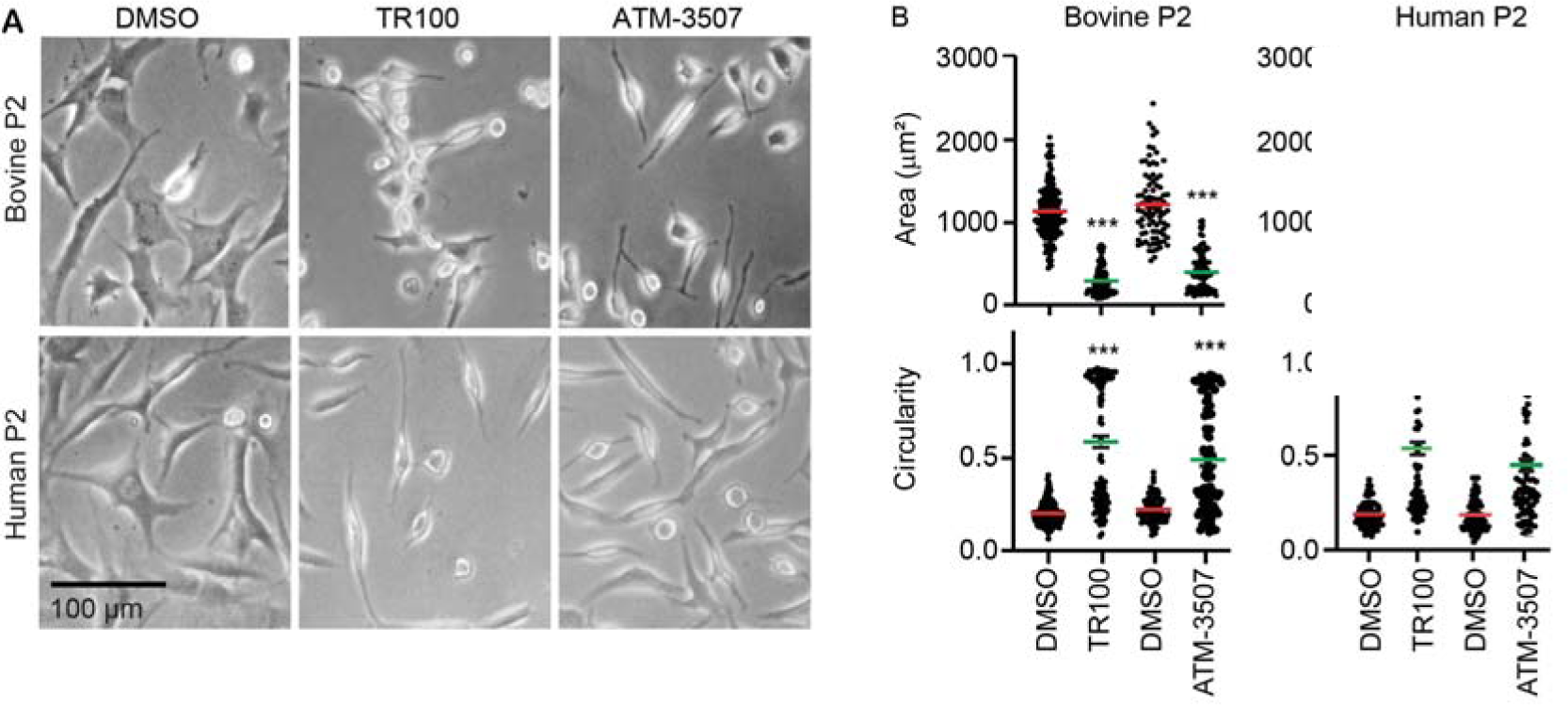
TPM3.1 inhibition leads to a change in cell morphology, actin reorganization, and cytoplasmic MRT localization. (A) Light microscopy images of bovine and human chondrocytes with TPM3.1 inhibitions, TR100 an ATM3507. (B) Quantification of light microscopy images showing that Tpm 3.1 inhibition decreases cell area an elongation in bovine (left) and human (right) P2 chondrocytes. ***, p < 0.001 as compared to DMSO control.

*TPM3.1 inhibition results in cortical F-actin distribution*

Exposure of passaged cells to TPM3.1 inhibitor abrogates the F-actin stress fibers leading to cortically organized F-actin (Figure 3A, B), resembling primary chondrocytes (Figure 2A). However, in contrast to primary chondrocytes, TPM3.1 is less enriched at the cortical regions of cells with diffuse staining throughout the cytoplasm. This is exemplified by less overlap in fluorescent intensity for TPM3.1 and F-actin in TPM3.1 inhibited passaged cells (Figure 3C) versus primary chondrocytes (Figure 2D).

**Figure 3.**
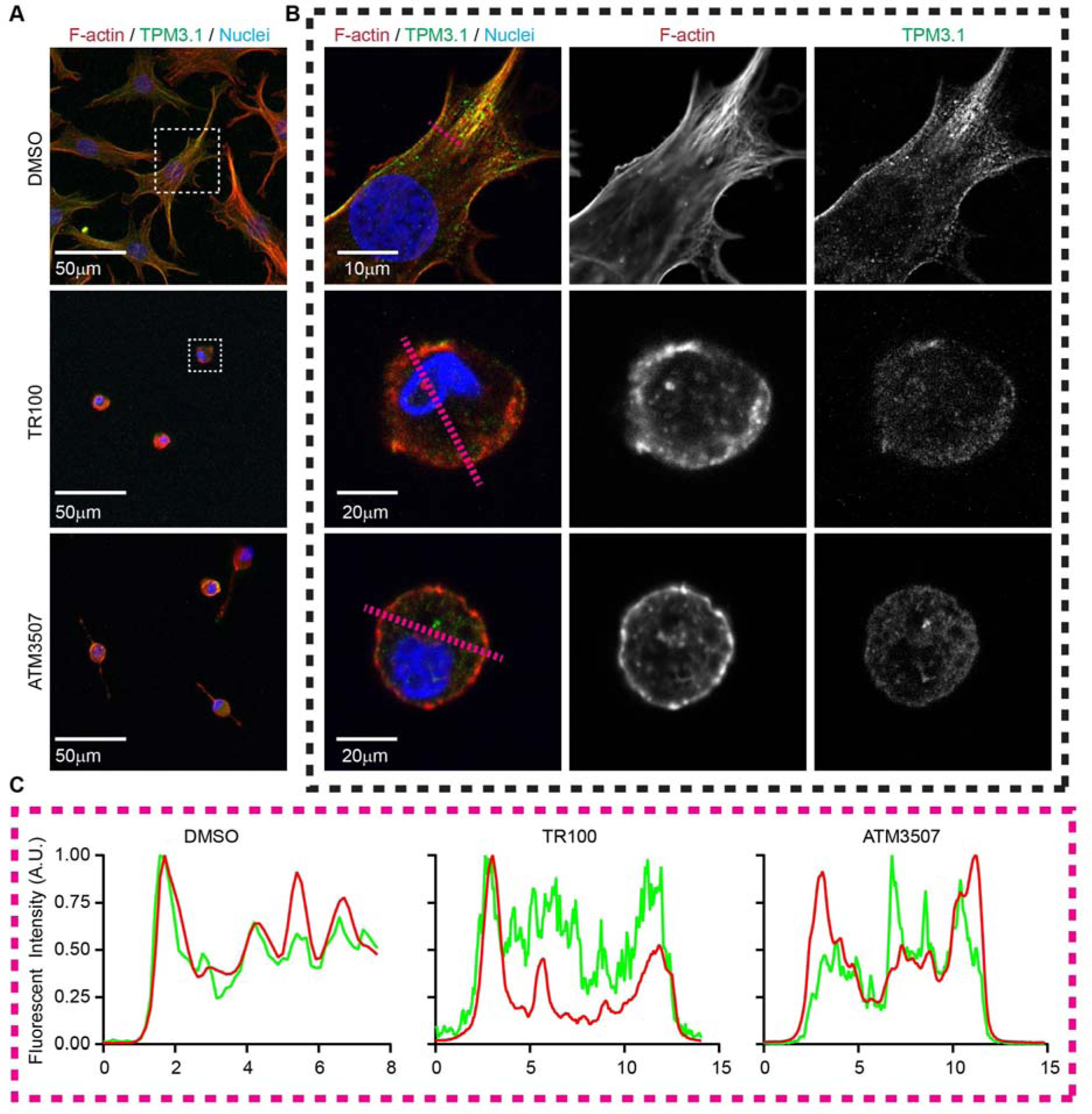
TPM3.1 inhibition ablates F-actin stress fibers and promotes cortical F-actin organization. (A) Lower magnification (20x objective) confocal microscopy images of bovine P2 cells treated with TPM3.1 inhibitors TR100 or ATM3507. (B) Higher magnification (63x objective) of regions in A (white dashed boxes) showing TR100 treatment in bovine cells leads to a cortical actin rearrangement. Cells stained for F-actin (Phalloidin; Red), TPM3.1 (Antibody 2G10.2; Green), and Nuclei (Hoechst; Blue). (D) Line scan analysis of lines (pink dashed) in panel B, demonstrating TR100 or ATM3507 treatment decreases colocalization between TPM3.1 with F-actin as compared to DMSO control. There is also less cortical TPM3.1 as compared to P0 cells (Figure 1A).

*Inhibition of Tpm 3.1 decreases nuclear MRTF and represses dedifferentiation molecule levels*

In our previous studies we found that abrogating F-actin stress fibers increases the proportion of G/F- actin, leading to nuclear export of MRTF causing repression of fibroblast matrix and contractile, as well as increases the expression of chondrogenic genes (Parreno et al., 2017a; Parreno et al., 2014; Parreno et al., 2017b). Therefore, we examined the effect of TPM3.1 inhibition by TR100 on F-actin organization and MRTF localization (Figure 4A, B). We determined that TPM3.1 inhibition by TR100, which abrogates F-actin stress fibers (Figure 3 and Figure 4A), also increases G/F-actin (Figure 4C, D) and reduces the proportion of nuclear MRTF (Figure 4B, E). TR100 decreases P2 cell *COL1*, *TNC*, α*SMA*, and *TAGLN* mRNA levels and increases *SOX9* mRNA levels. TR100 did not lead to upregulation of *ACAN* or *COL2* (data not shown).

**Figure 4.**
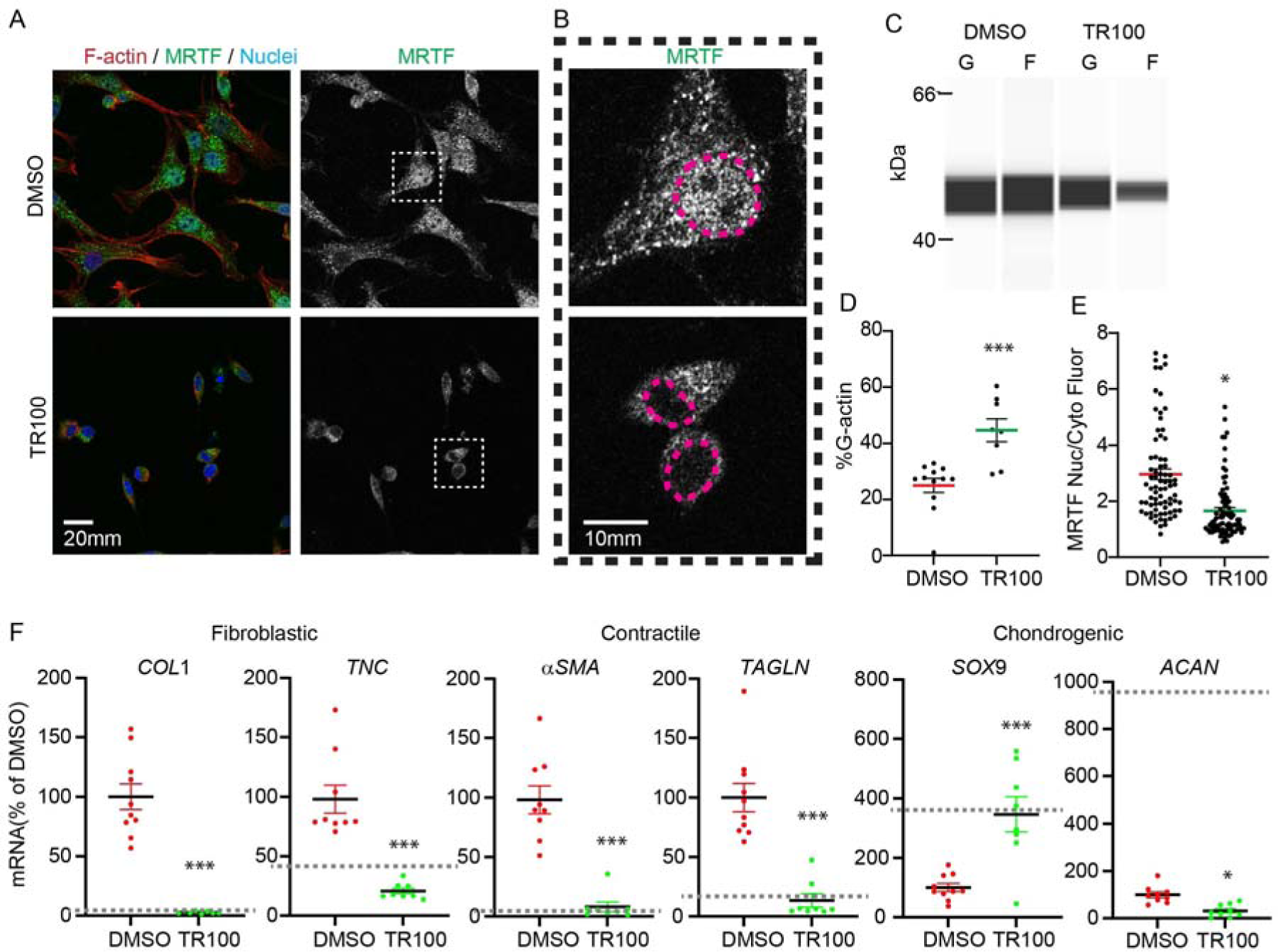
TPM3.1 inhibition in P2 chondrocytes by TR100 treatment partially redifferentiates P2 cells by reducing actin polymerization, nuclear MRTF, and dedifferentiated mRNA levels. (A) Lower magnification images (20x objective) of bovine P2 cells treated with TR100. Cells stained for F-actin (Phalloidin; Red), MRTF (Green), and Nuclei (Hoechst; Blue). (B) Higher magnification images (63x objective) of cells in A (white dashed boxes). Pink dashed tracings indicate nuclei borders. (C) Capillary Western blot (WES) immunoassay pseudoblot of Triton- fractionated samples for separation of G/F-actin. (D) Corresponding dot plot showing the increase in the proportion of G/F-actin in cells treated with TR100. (E) Corresponding dot plot of MRTF subcellular localization demonstrating a reduction in nuclear MRTF by TR100 treatment. (F) Relative real-time PCR of cells treated with TR100 demonstrating a reduction in fibroblastic and contractile genes as well as an increase in SOX (mRNA levels. *, p < 0.05; **, p < 0.01; ***, p < 0.001 as compared to DMSO treatment. Grey dashed lines in F represent primary (P0) bovine chondrocyte mRNA levels for genes based on our previous studies (Parreno et al., 2017a; Parreno et al., 2014; Parreno et al., 2017b).

The suppression of the dedifferentiated mRNA levels and upregulation of *SOX9* by TR100 treatment in bovine passaged chondrocytes, led us to examine if TR100 treatment would favorably regulate human P2 protein levels. TR100 treatment decreases COL1 and TAGLN expression as well as an increase in SOX9 expression (Figure 5).

**Figure 5.**
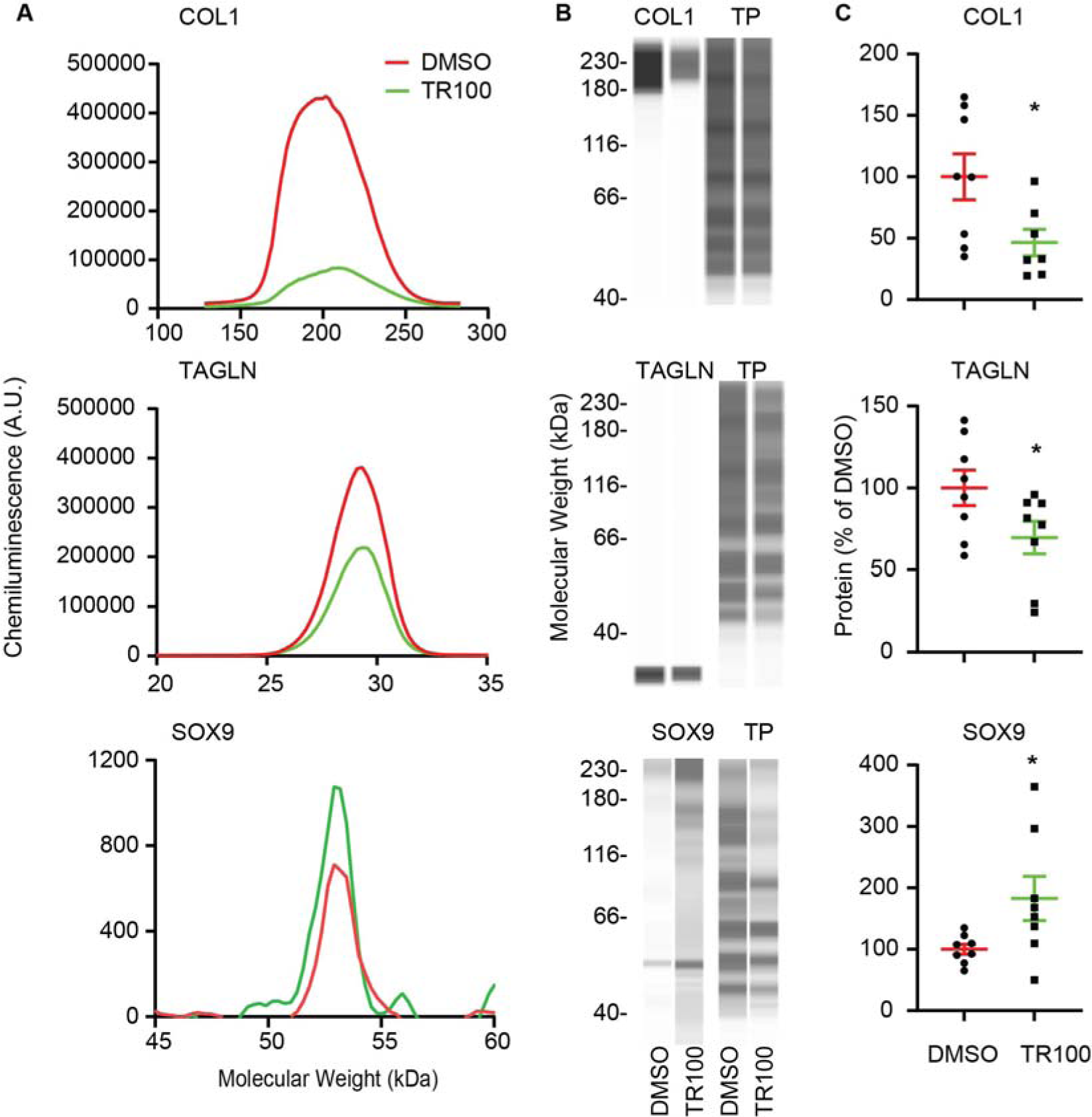
Modulation of select protein levels by TR100 treatment after 2 days of treatment. Representative WES capillary electrophoresis data showing representative (A) electropherograms and (B) pseudo-Western blots. Specific protein levels were normalized to total protein (TP) levels. (C) Corresponding dot plots of relative protein levels. Data demonstrates a decrease in fibroblast matrix (COL1) and contractile (TAGLN) protein levels as well as an increase in chondrogenic (SOX9) protein levels. *, p < 0.05 as compared to DMSO

Overall, our findings suggest that TPM3.1 is critical for maintenance of spread cellular morphology, F-actin stress fibers, nuclear MRTF localization, and the expression of dedifferentiated molecules.

### Inhibition of Cdc42 prevents cell spreading

Next, we sought to determine if targeting specific Rho-GTPases, which are known to regulate actin organization, may be another means to regulate the dedifferentiated phenotype in passaged chondrocytes by preventing respreading of P2 cells. To examine the role of RhoGTPases in cellular spreading, P2 cells were removed from culture vessels and re-seeded in the presence of the RAC, ROCK, and CDC42 inhibitors NCS23766, Y27632, and ML141 respectively. None of inhibitors affected the ability of P2 cells to attach as cells attached within 3 hours of seeding. Exposure of P2 cells to NSC23766 does not affect overall cell size or circularity (Figure 6A-C). Exposure of P2 cells to Y27632 slightly affected cell morphology with a reduction in average cell area (Figure 6A, C) as compared to DMSO treated controls. Y27632 treatment does not affect cell circularity (Figure 6A, C). ML141, however, has profound effects on morphology by inhibiting P2 cell spreading indicated by lower average cell size (Figure 6A, C). ML141 also increases average circularity (Figure 6A,C). Similar to treatment of P2 cells with TR100, there is a bimodal distribution of cell circularity consisting of rounded and elongated cell populations. Despite being elongated, cells were still smaller as compared to untreated control cells. Analysis of live cell, in incubator images, over time demonstrates that the effect of CDC42 inhibition on cell morphology is apparent by 12 and 24 hours (Figure 6A). Since CDC42 is a regulator P2 cell spreading, we examined the effects of CDC42 inhibition on the dedifferentiated phenotype for the remainder of our study. Similar to bovine P2 cells, we confirmed that ML141 also represses cell spreading and elongation in human P2 cells (Figure 6D-F).

**Figure 6.**
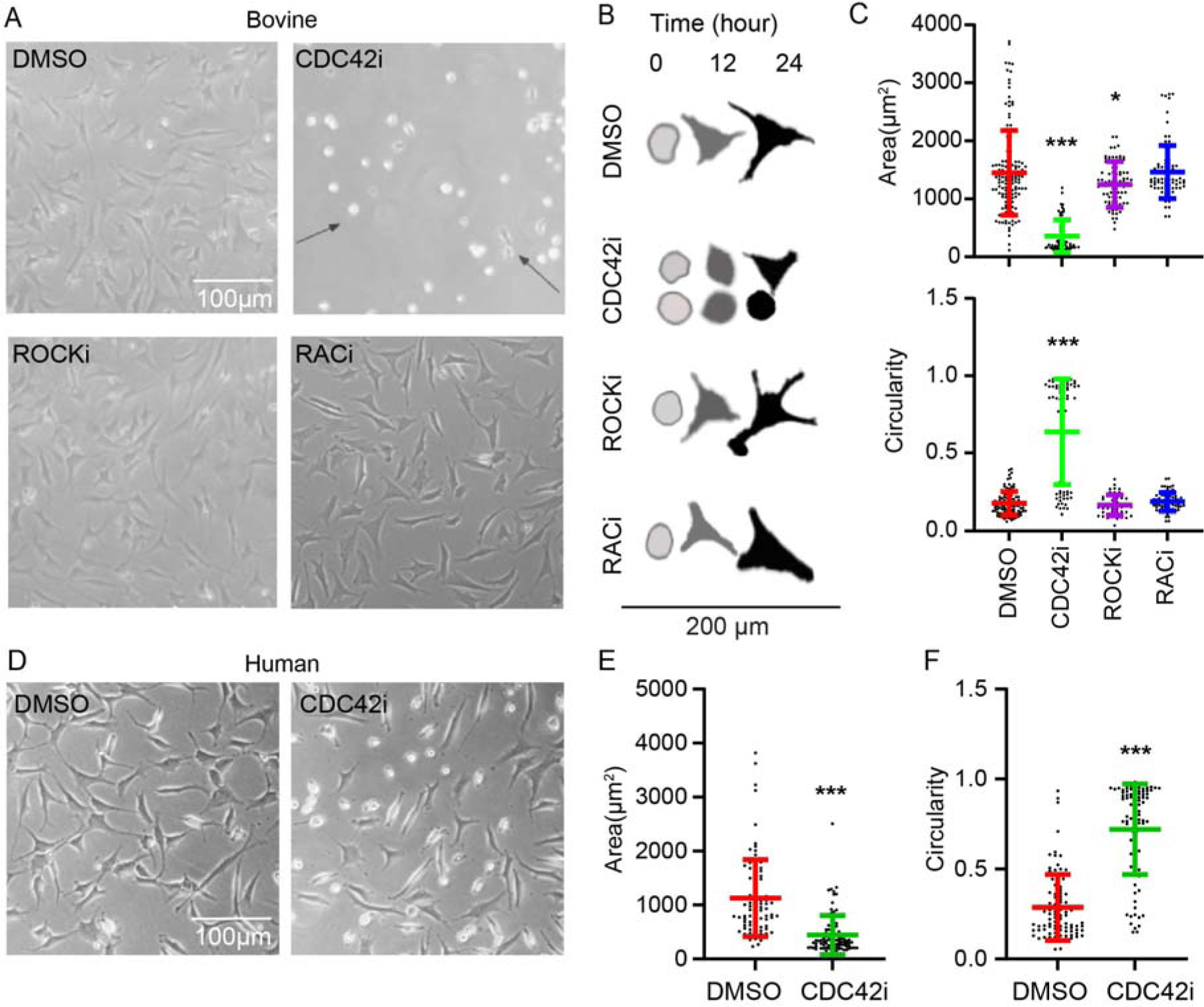
The effect of RhoGTPase inhibition on cell morphology of (A-C) bovine and (D-F) human P2 cells. (A) Light microscopy images of bovine chondrocytes exposed to the CDC42, ROCK, RAC inhibitors ML141 (CDC42i), Y27632 (ROCKi), and NSC23766 (RACi), respectively for 1 day. (B) Traces of cells from time-lapse in incubator imaging. (C) Quantification of light microscopy images showing that CDC42i decreases cell area and increases average cell circularity. Of note CDC42i led to bimodal distribution in cell circularity where a subpopulation of cells were smaller but remained elongated. Arrows in A and traces in B show both representative rounded and elongated cells. (D) Light microscopy of human P2 cells exposed to CDC42i for 1 day. (E) Quantification of light microscopy images showing that CDC42i decreases cell area and increases average cell circularity.

### CDC42 inhibition by ML141 increases G/F-actin and cytoplasmic MRTF

We next examined the effect of CDC42 inhibition on F-actin organization (Figure 7A) and MRTF localization (Figure 7B). Coinciding with alterations in morphology, ML141 treatment abrogated the formation of F-actin stress fibers (Figure 7A). As compared to DMSO treatment, ML141 treatment prevents F-actin stress fibers and rather leads to cortical F-actin organization (Figure 7A). In contrast to TR100 treatment (Figure 3), ML141 treatment results in TPM3.1 association with cortical F-actin (Figure 7A). ML141 increases the proportion of G/F-actin as compared to control cells (Figure 7C, D). Similar to TR100 treatment (Figure 4), ML141 treatment reduces nuclear MRTF (Figure 7B, E). This decreases fibroblastic matrix molecules, *COL1* and *TNC*, and contractile molecules, α*SMA* and *TAGLN* (Figure 7F). mRNA levels for ML141 treated P2 chondrocytes approached P0 mRNA levels for most of the genes. Additionally, *SOX9* showed a significant increase in mRNA expression, although not full recovery to P0 levels. ML141 did not increase the expression of *ACAN* or *COL2* (data not shown).

**Figure 7.**
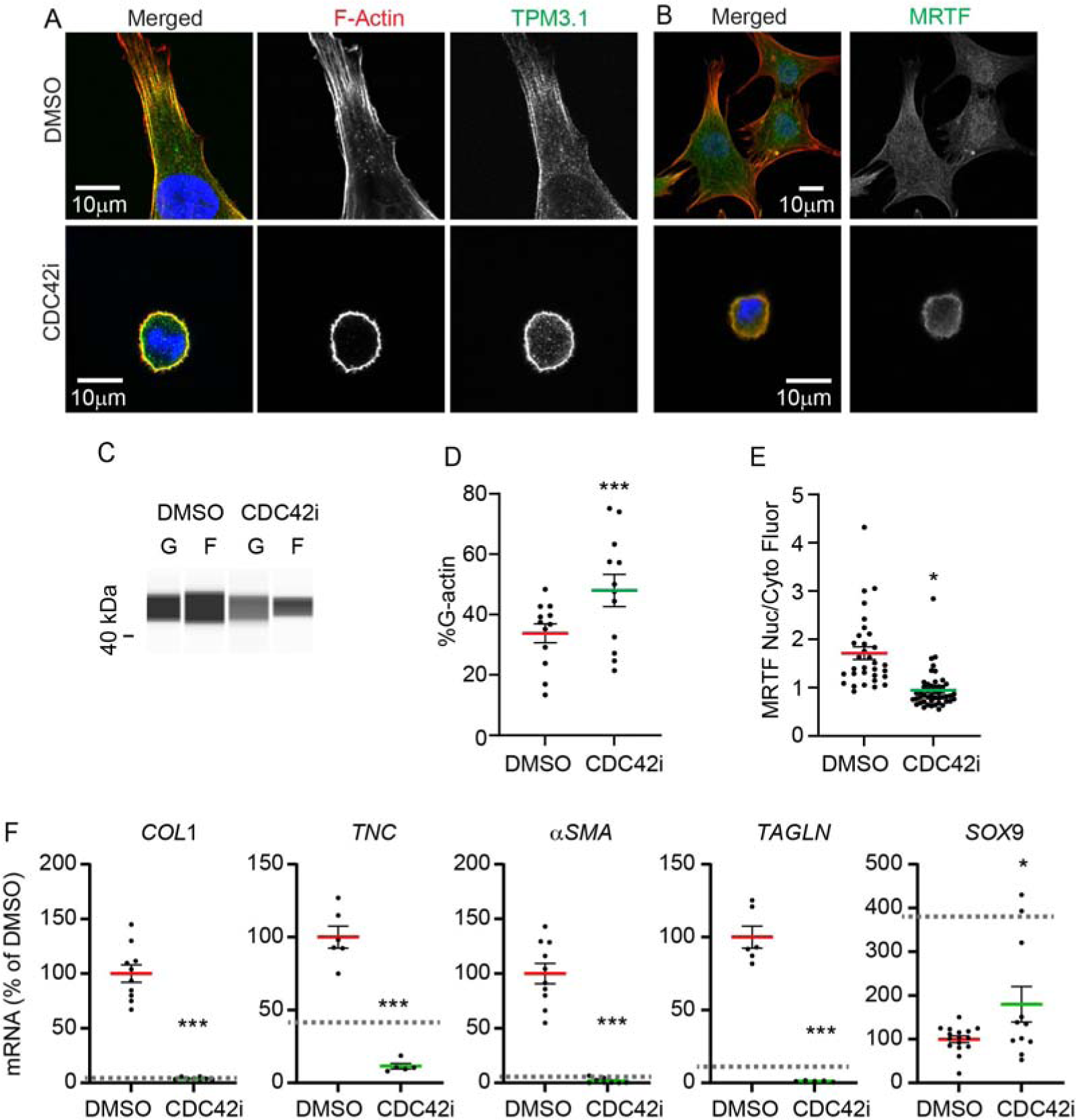
CDC42 inhibition by ML141 treatment leads to partial redifferentiation of bovine P2 cells by reducing F-actin stress fibers, nuclear MRTF, and dedifferentiated mRNA levels. (A, B) Images (63x objective) of cells treated with ML141 (CDC42i). Cells stained for F-actin (Phalloidin; Red), Tpm3.1 (Green; Panel A) or MRTF (Green; Panel B), and Nuclei (Hoechst; Blue). (C) Capillary Western blot (WES) immunoassay pseudoblot of Triton-fractionated samples for separation of G/F-actin. (D) Corresponding dot plot showing the increase in the proportion of G/F-actin in cells treated with ML141. (E) Corresponding dot plot of MRTF subcellular localization demonstrating a reduction in nuclear MRTF by ML141 treatment. (F) Relative real-time PCR of cells treated with TR100 demonstrating a reduction in fibroblastic and contractile genes as well as an increase in Sox9 mRNA levels. *, p < 0.05; **, p < 0.01; ***, p < 0.001 as compared to DMSO treatment. Grey dashed lines in F represent primary mRNA levels for genes based on our previous studies (Parreno et al., 2017a; Parreno et al., 2014; Parreno et al., 2017b).

Next, we examined the effect of ML141 treatment on human P2 mRNA and protein levels (Figure 8). Similar to bovine cells CDC42i reduces *COL1* and *TAGLN*. Unlike bovine P2 cells, ML141 did not upregulate *SOX9* mRNA levels. With regard to modulation of protein, ML141 significantly decreases COL1 and TAGLN protein levels (Figure 8B, C), similar to exposure of cells to TR100 (Figure 4). Despite no apparent differences in *SOX9* mRNA levels by ML141 treatment (Figure 8A), ML141 significantly increases chondrogenic Sox9 protein levels (Figure 8B, C).

**Figure 8.**
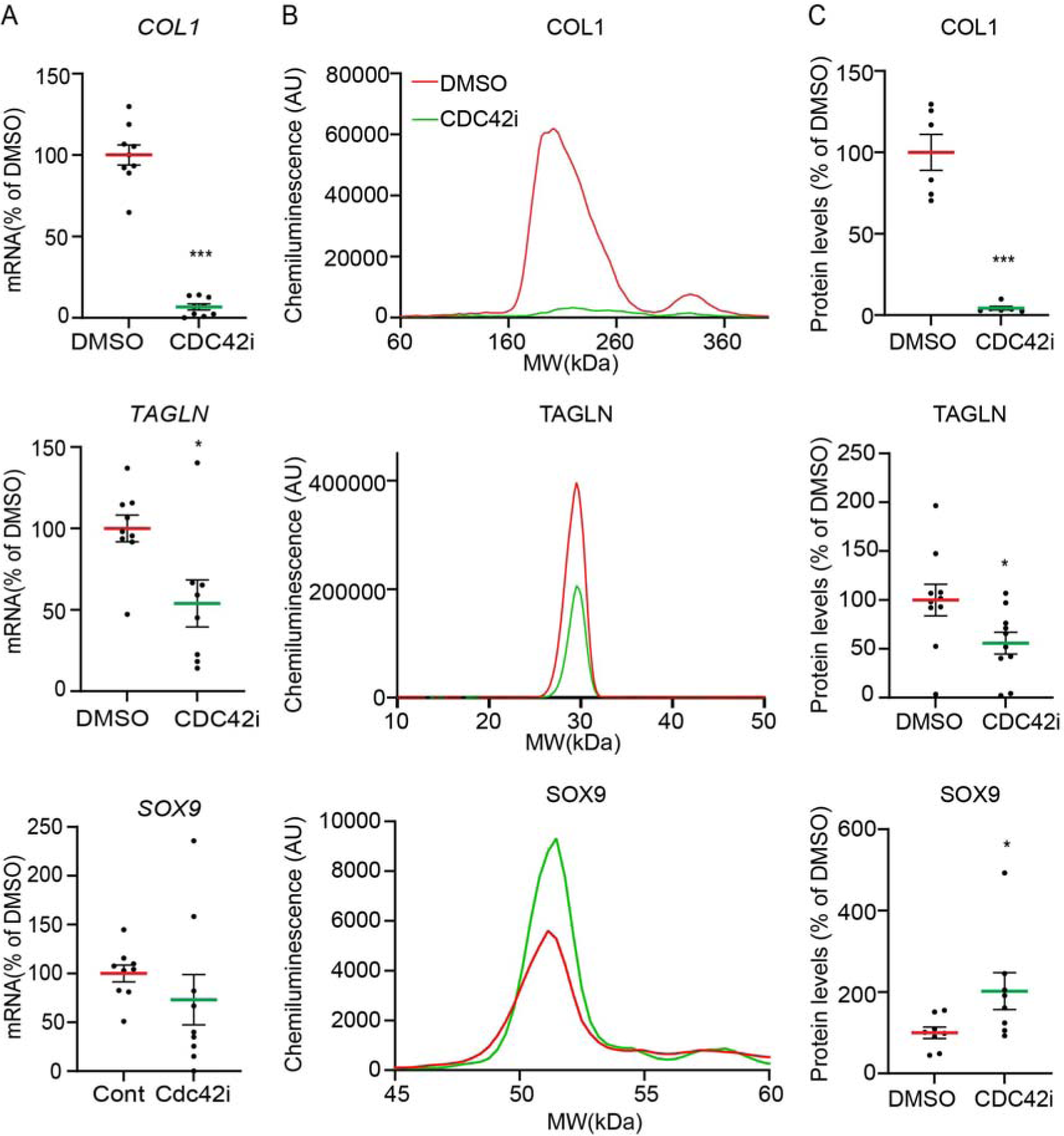
Modulation of select mRNA and protein levels in human P2 cells by ML141 treatment after 2 days of treatment. (A) Relative real-time PCR of cells treated with TR100 demonstrating a reduction in fibroblastic (*COL1*) and contractile (*TAGLN*) genes. (B) Representative WES capillary electropherograms. (C) Corresponding dot plots of relative protein levels after normalization to total protein demonstrating a decrease in fibroblast matrix (COL1) and contractile (TAGLN) protein levels as well as an increase in chondrogenic (SOX9) protein levels. *, p < 0.05; ***, p < 0.001 as compared to DMSO

## Discussion

This study provides new insights into the regulation of F-actin stress fibers in passaged cells. In support of our hypothesis, we determined inhibition of the F-actin network regulators, TPM3.1 or CDC42, led to similar findings in suppressing the dedifferentiated phenotype. TPM3.1 or CDC42 inhibition reorganized F-actin from stress fibers into cortical organization causing an increase in G/F-actin, reduction of nuclear MRTF, repression of dedifferentiated molecules, and an increase in SOX9 levels. Targeting TPM3.1 or CDC42 to repress stress fibers, is a means to suppress the dedifferentiated phenotype in monolayer expanded, passaged chondrocytes.

TPM3.1 is expressed in chondrocytes and associates with F-actin in stress fibers. To our knowledge, this is the first study to determine that TPM plays a critical role in regulating the chondrocyte phenotype. We found that TPM3.1 is critical in the maintenance of F-actin stress fibers in P2 chondrocytes which is in agreement with the regulation of stress fibers in lens epithelial cells (Parreno et al., 2020), tenocytes (Inguito et al., 2022), osteosarcoma (U2OS) cells (Gateva et al., 2017), human SK-N-BE(2) neuroblastoma cells, and neuronal B35 cells (Meiring et al., 2019; Pathan-Chhatbar et al., 2018; Tsolis et al., 2015; Zhao et al., 2020). TPM3.1 may stabilize of F-actin in stress fibers as well as regulate the binding of other actin-binding molecules onto F-actin. We previously have shown that Cofilin-1, which plays a role in severing F-actin, is expressed in passaged cells and promotes F-actin depolymerization (Parreno et al., 2017a). TPM3.1 binding can prevent the interaction of Cofilin onto F-actin (Bryce et al., 2003), therefore TPM3.1, in passaged cells, likely prevents F-actin depolymerization by Cofilin-1 dependent severing. Additionally, in other cell types, TPM3.1 can also enhance stress fibers by promoting the activation of non-muscle myosin (Bryce et al., 2003; Gateva et al., 2017; Pathan-Chhatbar et al., 2018; Tojkander et al., 2011). The role of TPM3.1 in the regulation of other critical actin binding proteins in chondrocytes is unknown and an area of future investigation. Furthermore, while we provide support that TPM3.1 is an important regulator of F-actin stress fibers, previous literature has indicated 5 other TPMs (1.6, 1.7, 2.1, 3.2, and 4.2) associate with stress fibers (Gateva et al., 2017; Tojkander et al., 2011). The expression and role of these other stress fiber associated TPMs have yet to be determined in chondrocytes.

In addition to TPM3.1, we demonstrate passaged cells require CDC42, but not RAC1 or ROCK, to spread out and organize F-actin into stress fibers, as well as to maintain the dedifferentiated phenotype. This contrasts with previous studies where RHOA as well as RAC1 regulates chondrogenesis (Haudenschild et al., 2010; Woods et al., 2005) (Wang and Beier, 2005; Woods et al., 2007). Additionally in contrast to our present study, a previous study demonstrated a negative relationship between CDC42 and F-actin stress fibers as reduction in CDC42 corresponded to an increase in stress fibers (Novakofski et al., 2012). A difference between previous studies and our present study, is we used passaged chondrocytes as opposed to chondroprogenitors or freshly isolated chondrocytes. This may suggest that differentiation state / cell-type may influence the regulation of cell phenoypte by specific RhoGTPases.

Our data demonstrated that TPM3.1 or CDC42 inhibition leads to a similar suppression of the dedifferentiated phenotype in P2 cells which we postulate both being mediated by the same actin-based regulation of MRTF. In addition to reducing stress fibers, TPM3.1 or CDC42 inhibition increase in G/F-actin and decrease in nuclear MRTF which we showed regulates fibroblastic matrix and contractile gene expression (Parreno et al., 2014; Parreno et al., 2017b). TPM3.1 and CDC42 inhibition has similar effects as latrunculin on P2 cells (Parreno et al., 2014; Parreno et al., 2017b); exposure of P2 cells to latrunculin also increases G/F-actin, decreases nuclear MRTF, reduces dedifferentiated (fibroblast matrix, contractile) molecule expression and increases SOX9. Another similarity is that inhibition of TPM3.1, CDC42, or treatment with Latrunculin did not lead to full redifferentiation of passaged cells as *COL2* mRNA levels were not recovered by treatments. Cortical F-actin is thought to be critical in the maintenance of the differentiated chondrocyte phenotype including *COL2* expression (Park et al., 2008). Unlike latrunculin which prevents cortical F-actin formation, TPM3.1 or CDC42 inhibition enables the formation of cortical F-actin. However, despite the reorganization into cortical F-actin, *COL2* mRNA levels are not recovered by either TPM3.1 or CDC42 inhibition. This suggests that other factors other than cortical F-actin are also required to recover COL2 expression. Nevertheless, the regulation of P2 F-actin through TPM3.1 or CDC42 inhibition can prime redifferentiation and it has been shown the formation of cortical F-actin may be an initial step in the redifferentiation process (Parreno et al., 2018).

While we show that both TPM3.1 and CDC42 are regulators of F-actin networks, the relationship between CDC42, TPM3.1, and specific F-actin network organizations remains to be determined. Data from another study has led to the suggestion that RhoA promotes F-actin formation and the association of TPM3.1 along cortical F-actin (Wang et al., 2023). Interestingly, in the present study we found that inhibition of CDC42 abrogates F-actin stress fibers and promotes cortical F-actin, which TPM3.1 associates with. It comes into question whether inhibition of CDC42 may lead to greater proportion of RHOA activity to promote stabilization by specific binding of TPM3.1 to cortical F-actin. An interesting difference between CDC42 and TPM3.1 inhibition, is that TR100 treatment did not lead to the association of TPM3.1 along cortical F-actin. The consequence on the lack of TPM3.1 along cortical F-actin in chondrocytes is unknown. One postulation is that a lack of TPM3.1 along cortical F-actin may lead to reduced cortical F-actin stability and susceptibility of cortical F-actin to depolymerization. As loss of cortical F-actin has recently been shown to be associated with cell death during disease pathology in Osteoarthritis (Chan et al., 2023), careful consideration on which actin mechanisms to target to prime redifferentiation for cell-based therapies and/or bioengineering is required. Targeting CDC42 may be a means to achieve cortical F-actin networks with TPM3.1 association that resemble the primary chondrocyte state.

In summary, this work shows two potential targets to suppress the dedifferentiated phenotype and prime redifferentiation in passaged chondrocytes via targeting regulators of F-actin. This understanding may aid in the development of new strategies to improve cell-based therapies and/or the generation of bioengineered cartilage.

## Acknowledgements

The research reported in this project was supported by the Delaware Center for Musculoskeletal Research from the National Institutes of Health’s National Institute of General Medical Sciences under grant – NIGMS (P20 GM139760). This publication was made possible by the Delaware INBRE program, supported by a grant from the National Institute of General Medical Sciences – NIGMS (P20 GM103446) from the National Institutes of Health and the State of Delaware.

## Supplementary Data

**Figure S1.**
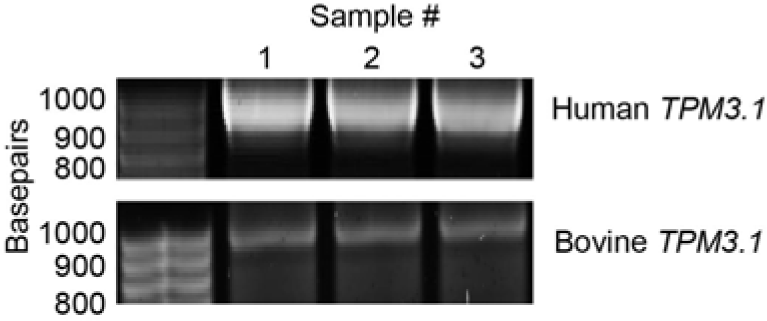
Semi quantitative PCR for *TPM3*.1 in human (top) and bovine (bottom) passaged cells. Bands were cut from gels and verified to be *TPM3*.1 by Sanger Sequencing.

